# Asymmetric dispersal alters extinction dynamics in a two-patch metapopulation under an Allee effect

**DOI:** 10.64898/2026.11.23.765930

**Authors:** Margaret E. Swift, King-Yeung Lam, Leah Shaw, Junping Shi

## Abstract

This paper investigates a two-patch metapopulation model with asymmetric dispersal, motivated by the directional transport of organisms between populations by aquatic currents. We consider a system of ordinary differential equations in which dispersal is population-conserving but directionally asymmetric, and each patch is subject to a strong Allee effect: Populations below a critical thresh-old decline toward extinction, whereas those above the threshold persist and approach their carrying capacities. We analytically prove the convergence of solutions to equilibria, and identify parameter regimes in which positive equilibria necessarily exist. We then numerically explore the one- and two-parameter steady-state bifurcation structures with respect to combinations of asymmetric dispersal rates and the Allee threshold. In particular, for sufficiently high Allee thresholds, we identify broad parameter regions in which the extinction equilibrium is the unique equilibrium of the system. These extinction regions arise from the interplay between asymmetric dispersal and the Allee effect and, notably, are absent in the corresponding population-conserving dispersal model without an Allee effect. Our results demonstrate that, through ‘rescue’ effects and emergent extinction regimes, directional dispersal can fundamentally help or hinder population persistence in systems subject to the Allee effect, highlighting important implications for the conservation and restoration of endangered species experiencing asymmetric dispersal.

**MSC 2020:** 92D25, 34C23, 34D05, 34D45, 92D40

## 1 Introduction

Asymmetric, gradient-driven ecological processes are common in the natural world, from the directed transport of oyster spat along the water column (Zang et al. 2026) to seed dispersal in anemochoric plants (Liang et al. 2026). These processes can be heavily modified by positive density-dependent processes such as the Allee effect, in which population decreases when under a characteristic critical density known as the Allee threshold (Allee 1931). Understanding how asymmetric dispersal can augment the Allee effect is critical for management and recovery of low-density metapopulations, as movement between high- and low-density patches could either rescue or deplete neighboring populations of the same species.

Asymmetric dispersal can be driven by environmental and behavioral mechanisms. Organisms that rely on the environment to move are bound by the direction of natural currents such as aquatic currents or wind current (Vuilleumier et al. 2010). In river and stream networks, organisms and their larvae, macroinvertebrates, and plant seeds are swept downriver easily, while upstream movement requires active, high-energy swimming. For example, in populations of freshwater pearl mussel (*Margaritifera laevis*) unidirectional river flow produces strongly downstream-biased dispersal, while upstream movement is limited to a short larval stage associated with host fish (Terui et al. 2014). The resulting connectivity between populations can therefore be strongly directional (Cowen and Sponaugle 2009).

The Allee effect has its greatest impact on populations at low density, which are often members of endangered species or subject to concerted recovery efforts. For example, large-scale melting of solid sea ice may limit mating opportunities for vulnerable polar bear (*Ursus maritimus*) populations below a critical threshold (Molnar et al. 2008); human-induced habitat fragmentation may constrain cooperative pack hunting in the endangered African wild dog (*Lycaon pictus*) (Courchamp et al. 2000); and declining colony sizes limit anti-predator vigilance in the critically endangered Vancouver Island marmot (*Marmota vancouverensis*) (Brashares et al. 2010). The Allee effects is not limited to the animal kingdom; for instance, human harvesting of American ginseng (*Panax quinquefolius*) has fragmented populations into patches that may be too small to persist (Hackney and McGraw 2001), and plants that depend on animal pollinators suffer strong Allee effects at low population density (Hale et al. 2022). Often, anthropogenic disturbance reduces population densities, contributing to higher instances of the Allee effect and extinction risk (Courchamp et al. 2008).

Mathematical exploration of the Allee effect starts by examining the simpler logistic growth model, which accounts only for negative density-dependent effects, such as crowding and competition for limited resources (Seidl and Tisdell 1999). In the continuous-time logistic model, every positive solution converges to the carrying capacity as time tends to infinity. Mathematically, the model has a unique positive equilibrium, corresponding to the carrying capacity, which is globally asymptotically stable, while the trivial extinction equilibrium is unstable. This fundamental dynamical property extends to spatially structured logistic growth models, including coupled patch systems and reaction-diffusion models (Amarasekare 1998; Cantrell and Cosner 2003; Cosner 1996; Du and Li 2014; Li and Shuai 2010; Lu and Takeuchi 1993). In coupled patch models, the connectivity of the network determines how individual habitat patches are linked through species migration. Greater connectivity can reduce the risk of local extinction by facilitating the rescue effect, whereby immigration from neighboring patches sustains declining local populations, and can also promote the recolonization of previously occupied patches or the colonization of previously unoccupied patches (Brown and Kodric-Brown 1977; Eriksson et al. 2014). In particular, the existence and uniqueness of a globally asymptotically stable positive equilibrium in continuous-time patch models with logistic growth provides a mathematical foundation for population persistence and the effectiveness of dispersal-mediated rescue and recolonization.

Contrastingly, increasing the growth rate in an isolated discrete-time logistic system can lead to periodic oscillations and, eventually, chaotic dynamics (Amarasekare 1998; Doebeli 1995; Hastings 1993). By contrast, dispersal and coupling among non-isolated multi-patch communities can play an important role in stabilizing otherwise periodic or chaotic population dynamics. A variety of coupling mechanisms have been shown to promote stabilization or coexistence, including migration between source and sink habitats (Holt 1985); low-rate passive symmetric (Hastings 1993) and asymmetric (Doebeli 1995) dispersal between two coupled patches; asynchronous reproduction and dispersal (Doebeli 1995); weak coupling in the absence of external noise (Ruxton 1994); and mortality during migration between patches (Amarasekare 1998). Collectively, these studies demonstrate that spatial coupling can substantially suppress the periodic and chaotic dynamics that arise in isolated logistic populations.

Discrete-time population models with Allee effects can exhibit four distinct dynamical behaviors: persistence, bistability, extinction, and essential extinction, in which almost all initial population densities lead to extinction. Rapidly growing populations may persist even under high predation pressure, but experience essential extinction when predation is reduced (Schreiber 2003). In two-patch discrete-time models with Allee effects, multiple coexisting attractors can arise. A rescue effect may occur under weak coupling when one subpopulation is above the Allee threshold while the other is below it, or under strong coupling when both subpopulations remain above the Allee threshold and their local dynamics are out of phase (Vortkamp et al. 2020). Further-more, the strength of connectivity between the two patches can significantly influence the asymptotic total population size (Grumbach and Hilker 2025).

Multi-patch dynamics have also been investigated for continuous-time metapopulation models incorporating the Allee effect. In contrast to logistic systems, the dynamics of Allee-effect models can depend sensitively on the number of patches and the structure of dispersal among them. For example, dispersal coupled with competition can sustain persistent heterogeneity between two populations subject to an Allee effect (Gyllenberg et al. 1999). Dispersal can also enable patches with populations below their local Allee thresholds to persist through immigration from neighboring patches with more robust populations, thereby producing a rescue effect (Amarasekare 1998; Gyllenberg et al. 1999; Kang and Lanchier 2011). In particular, when dispersal is symmetric and population-conserving, this rescue mechanism can lead to the existence of positive equilibria (Kang and Lanchier 2011). Focusing on a two-patch system with heterogeneous Allee thresholds, Kang and Lanchier considered lossless, symmetric, and density-independent dispersal and showed that the system can possess between two and eight positive equilibria, depending on the relative values of the Allee thresholds in the two patches (Kang and Lanchier 2011). In contrast, when dispersal involves mortality, the rescue effect can be weakened or eliminated: Sufficiently high Allee thresholds combined with moderate-to-low survival rates during dispersal can lead to global extinction.

Here, we study a two-patch coupled system of ordinary differential equations experiencing asymmetric migration and an Allee effect growth rate. First, we introduce our model and its derivation, then show the convergence of solution trajectories to equilibria (by defining a Lyapunov function for the system), which implies absence of closed orbits. We then we prove the existence of positive equilibria in certain parameter ranges, and also prove the nonexistence of positive equilibria in other ranges. These analytic findings are complemented by numerical bifurcation analysis to reveal a more complete picture of global dynamics of the system for all parameter ranges. Lastly, we conclude with a discussion of extinction parameter zones in which no initial combination of populations can persist, and an emergent population rescue effect achieved through asymmetric dispersal. These global extinction and rescue effect zones are a significant finding for conservation efforts in species with directed dispersal and positive density dependence under a critical density threshold.

## 2 Mathematical Model

Throughout the paper, ℝ is the set of all real numbers, ℝ^+^ = (0, +*∞*) 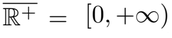 and. To build our model, we consider two coupled populations that are subject to the Allee effect and asymmetric dispersal. A prototypical population model of the Allee effect growth is of the form

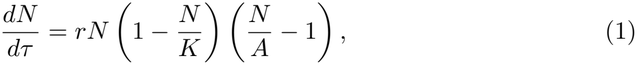

where *τ* is the time, *N* (*τ*) is the population of a biological species, *r >* 0 is the growth parameter, *K >* 0 is the carrying capacity, and *A* is the Allee threshold satisfying 0 *< A < K* (Sato 2009; Wang et al. 2011, 2019). To include density-independent dispersal in a two-patch model, we add an immigration term and subtract an emigration term from each patch:

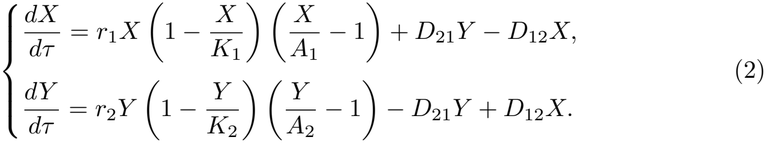

The following parameters apply to populations *X*(*τ*) and *Y* (*τ*), respectively: *r*_1_ and *r*_2_ are the growth parameters, *K*_1_ and *K*_2_ the carrying capacities, and *A*_1_ and *A*_2_ the Allee thresholds. The migration rate from patch *Y* to *X* is given by *D*_21_, while the one for migration from *X* to *Y* is *D*_12_, where *D*_12_, *D*_21_ ≥ 0. Dispersed population is completely conserved by balancing the migration terms for both patches; no organisms die while traveling between patches.

For this paper, we assume the population growth in the two patches is identical, and we focus on the effect of asymmetric dispersal on the two connected identical populations. Let *K*_1_ = *K*_2_ = *K, A*_1_ = *A*_2_ = *A* and *r*_1_ = *r*_2_ = *r*, and we make the following change of variables to non-dimensionalize Equation (2):

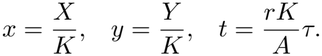

Then the model (2) is reduced to the following set of equations

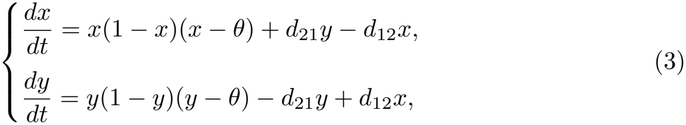

where

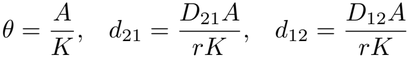

are the scaled Allee threshold and dispersal rates. Note that, due to our non-dimensionalization, *θ* ∈ (0, 1) and *d*_12_, *d*_21_ *≥* 0.

## 3 Analytic Results

### 3.1 Lyapunov function

Here we consider a general two-patch model with possibly different local dynamics as follows:

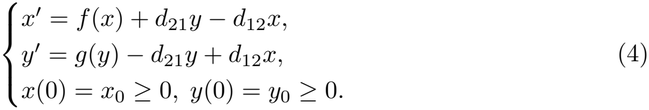

Here we assume that 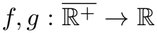 are smooth functions, and *d*_12_, *d*_21_ *≥* 0. Define a function 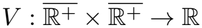 by

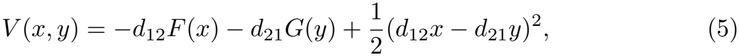

where

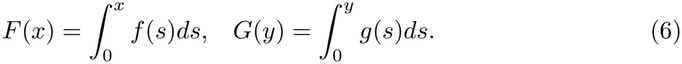

Then we can observe that (4) is equivalent to

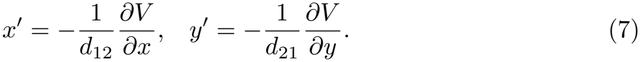

Hence (4) is almost a gradient system and *V* (*x, y*) serves as a Lyapunov function of the system (4).

We have a general result on the asymptotic behavior of solutions of (4) as follows.

#### Theorem 3.1.

*Suppose that* 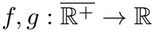 *are continuously differentiable functions, and d*_12_, *d*_21_ *>* 0. *In addition, we assume that*

i. *f* (0) *≥* 0 *and g*(0) *≥* 0;
ii. *Let V* (*x, y*) *be defined as in* (5). *Then there exists C* ∈ ℝ *such that V* (*x, y*) *≥ C for all x, y ≥* 0, *and the set S*_*a*_ = *{*(*x, y*) : *x, y ≥* 0, *V* (*x, y*) *≤ a} is bounded for any a* ≥*C;*
iii. *Each equilibrium point of* (4) *is isolated*.

*Then*

a. 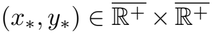 *is an equilibrium point if and only if* (*x*_*_, *y*_*_) *is a critical point of V, that is* ∇*V* (*x*_*_, *y*_*_) = (0, 0).
b. *For any* 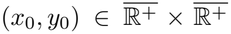, (4) *has a unique solution* (*x*(*t*), *y*(*t*)) *defined for t* ∈ (0, *∞*) *such that x*(*t*) *≥* 0 *and y*(*t*) *≥* 0 *for t* ∈ (0, *∞*). *Moreover, there exists an equilibrium point* 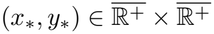 *such that* 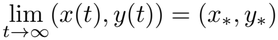 .
c. *System* (4) *has no periodic orbits*.

*Proof* Part (a) is obvious from (7), and part (c) follows from part (b). So we only need to prove part (b). If 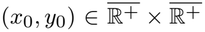, then a local solution (*x*(*t*), *y*(*t*)) exists following the standard existence and uniqueness result for ordinary differential equations. Since *f* (0) *≥* 0, then *x*^*’*^ = *f* (0) + *d*_21_*y ≥* 0 if *x*(*t*_0_) = 0 and *y*(*t*_0_) *≥* 0 for some *t*_0_ *≥* 0, and similarly *y*^*’*^ = *g*(0) + *d*_12_*x ≥* 0 if *x*(*t*_0_) *≥* 0 and *y*(*t*_0_) = 0 for some *t*_0_ *≥* 0. Hence the positive quadrant is positively invariant.

To show *V* is a Lyapunov function, we calculate that

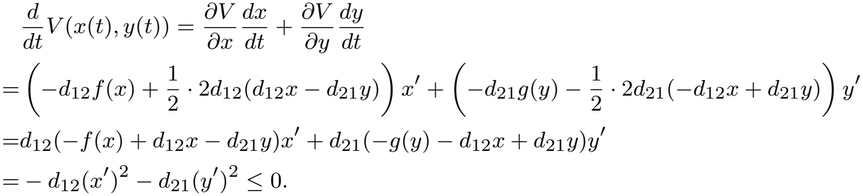

From (ii), any orbit (*x*(*t*), *y*(*t*)) of (5) is bounded thus exists for all *t >* 0. Now applying LaSalle’s Invariance Principle (LaSalle 1960, Theorem 2), we conclude that every solution (*x*(*t*), *y*(*t*)) starting from 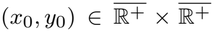 approaches *M* as *t → ∞*, where *M* is the largest invariant subset of 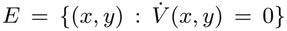 which is the set of equilibria from part (a). Finally, as all elements of *E* are isolated, then (*x*(*t*), *y*(*t*)) must converge to a single equilibrium point as *t → ∞*. □

The construction of Lyapunov function in (5) is clearly motivated by the Lyapunov function

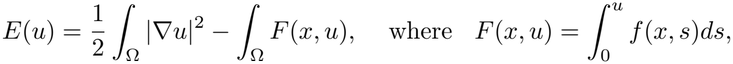

for the reaction diffusion equation

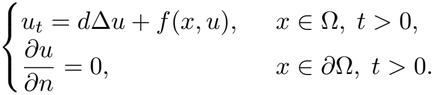

However, there is no general way to extend this construction to *n*-patch models with *n*≥ 3 unless the dispersal is symmetric, *d*_*ij*_ = *d*_*ji*_.

We remark that for the general system (4) or (7), the local stability of an equilibrium (*x*_*_, *y*_*_) is related to the Lyapunov function *V* . Since the Jacobian matrix of (7) is

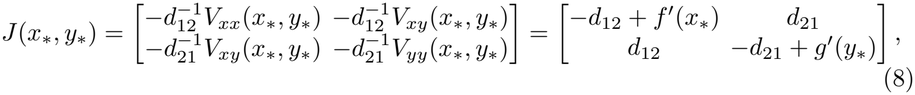

then (*x*_*_, *y*_*_) is a sink for (4) if it is a local minimum point of *V* ; (*x*_*_, *y*_*_) is a source for (4) if it is a local maximum point of *V* ; and it is a saddle point for (4) if it is a saddle point of *V* .

Now to show that Theorem 3.1 can be applied to (3), we prove the following proposition.

#### Proposition 3.2.

*Suppose that d*_12_, *d*_21_ *>* 0, *and f* (*x*) = *g*(*x*) = *x*(1 *− x*)(*x − θ*) *for some θ* ∈ (0, 1) *in* (4). *Then the conditions* (i)*-*(iii) *in Theorem 3.1 are satisfied. Hence system* (3) *has no periodic orbits, and for any* 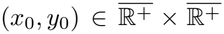, (3) *has a unique solution* (*x*(*t*), *y*(*t*)) *defined for t* ∈ (0, *∞*) *such that x*(*t*) *≥* 0 *and y*(*t*) *≥* 0 *for t* (0,), *and there exists an equilibrium point* 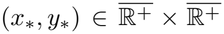 *such that* 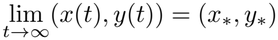.

*Proof* The condition (i) is obvious as *f* (0) = *g*(0) = 0. For (ii), we have

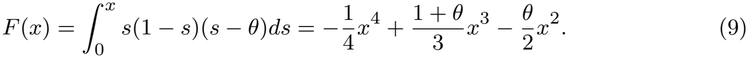

Thus we have

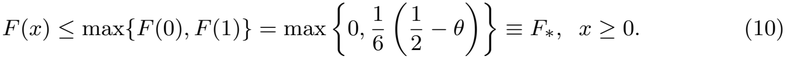

Then for any *x, y ≥* 0,

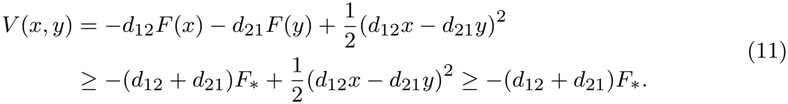

On the other hand, *V* (*x, y*) is of the form

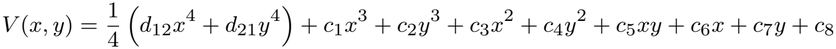

where *c*_*ϵ*_are constants. By applying various forms of Young’s inequalities,

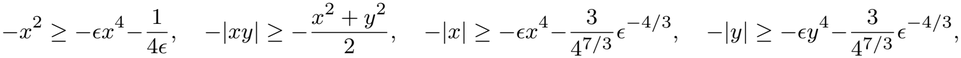

we may estimate *V* (*x, y*) from below by

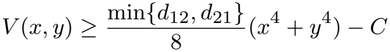

where *C >* 0 is a constant depending on *c*_*ϵ*_. Hence, for each *a >* 0

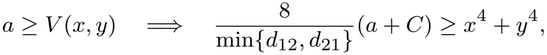

This proves (ii).

For (iii), we notice that the equilibrium equations of (3) are

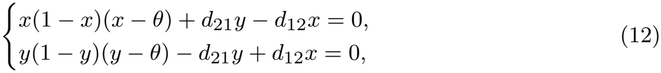

which can be reduced to a degree 9 polynomial equation of *x*. Hence (3) has at most 9 equilibria, and each of them must be isolated. □

### 3.2 Existence and stability of positive equilibria

Proposition 3.2 shows that the asymptotic dynamics of (3) is completely determined by the set of equilibria of (3). System (3) always has a trivial equilibrium (0, 0). The stability of the trivial state can be determined through the Jacobian matrix of (3) at (0, 0). Indeed from (8) we have

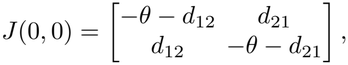

and the eigenvalues of *J* (0, 0) can be calculated as

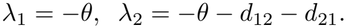

Since *θ* ∈ (0, 1) and *d*_12_, *d*_21_ ≥ 0, then both eigenvalues are negative. Therefore the trivial state (0, 0) is a sink and it is locally asymptotically stable.

Next we show that when 0 *< θ <* 1/2, then (3) always possesses at least two positive equilibria.

#### Theorem 3.3.

*Suppose that* 0 *< θ <* 1/2. *Then for every d*_12_, *d*_21_ *>* 0, *the system (3) has at least two positive equilibria*.

The proof of Theorem 3.3 is given at the end of this section. Note that for the single patch model

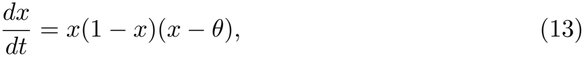

*V*_1_(*x*) = −*F* (*x*) is a Lyapunov function, and *x* = 0 and *x* = 1 are the two local minimum points of *V*_1_(*x*). The condition 0 *< θ <* 1/2 makes *x* = 1 the global minimum of *V*_1_(*x*) as 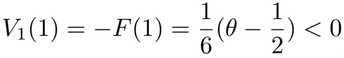 .

When the Allee threshold *θ* satisfies 1/2 *≤ θ <* 1, the trivial equilibrium *x* = 0 is the global minimum of *V*_1_(*x*). Indeed, in this case, it is possible that the extinction state (0, 0) is the only nonnegative equilibrium of (3) (see next subsection). In the following we show that system (3) possesses at least one positive equilibrium when *d*_12_ or *d*_21_ is small.

#### Theorem 3.4.

*Suppose that* 1*/*2 *≤ θ <* 1 *and*

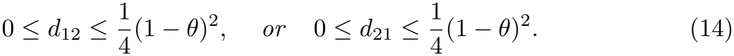

*Then* (3) *has at least one positive equilibrium*.

*Proof* Suppose that 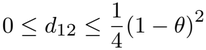 . We consider an auxiliary scalar equation

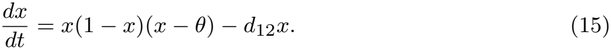

Then it is easy to see that besides the trivial equilibrium *x* = 0, (15) has two positive equilibria given by

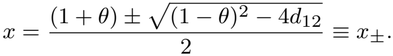

Now consider a solution (*x*(*t*), *y*(*t*)) of (3) with initial condition (*x*(0), *y*(0)) = (*x*_+_ +1, *y*_0_) with arbitrary *y*_0_ *≥* 0. From Theorem 3.1, the solution (*x*(*t*), *y*(*t*)) is positive when *t >* 0. Hence

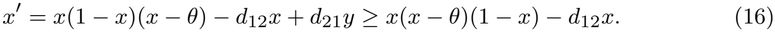

From the dynamics of (15), we know that *x*(*t*) *≥ x*_+_ for all *t >* 0. From Theorem 3.1, the solution (*x*(*t*), *y*(*t*)) tends to a nonnegative equilibrium (*x*_*_, *y*_*_) of (3) with *x*_*_ *≥ x*_+_, and *y*_*_ *≥* 0. If *y*_*_ = 0, then *x*_*_ = 0 from the second equation of (3). Hence *y*_*_ *>* 0 and (*x*_*_, *y*_*_) is a positive equilibrium. The proof for the case that 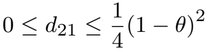 is similar. □

When the conditions on *d*_12_, *d*_21_, *θ* in Theorem 3.3 and 3.4 are satisfied, a conditional persistence holds for (3): when the initial condition is large, then the population in both patches will sustain; and when the initial condition is small, then the population in both patches will become extinct as the trivial state (0, 0) is always locally asymptotically stable. In Section 4 we will numerically show that generically (3) has 2, 4, 6 or 8 positive equilibria depending on the parameter values of *d*_12_, *d*_21_, *θ*, and it is also possible that (3) has no positive equilibrium.

Now we give the proof of Theorem 3.3. We first transform (*x, y*) = (*x, τx*) and note that (*x, y*) is a positive equilibrium of (12) if and only if

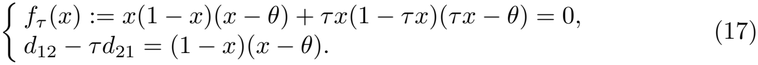

In the following we shall consider (without loss of generality) the case *d*_12_ *≥ d*_21_, where it is intuitive to see that if (*u, v*) = (*x, τx*) is a positive equilibrium, then *τ ≥* 1. We first show the following lemma on the properties of the function *f*_*τ*_ (*x*) for different values of *θ*.

#### Lemma 3.5.

*Let f*_*τ*_ (*x*) *be given as in* (17).

a. *Let θ* ∈ (0, 1*/*2). *Then for each τ* ≥ 1, *there exist two distinct positive numbers x*^*^(*τ*), *x*^**^(*τ*) *(depending continuously on τ) such that f*_*τ*_ (*x*^*^(*τ*)) = *f*_*τ*_ (*x*^**^(*τ*)) = 0 *and*

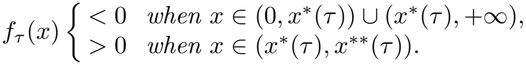

*Moreover, x*^*^(1) = *θ and x*^**^(1) = 1.

a. *Let θ* ∈ (1*/*2, 1). *Then there exists* 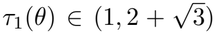 *and* 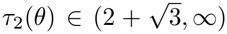, *such that for all τ* ∈ (*τ*_1_(*θ*), *τ*_2_(*θ*)), *f*_*τ*_ (*x*) *<* 0 *for all x >* 0. *Furthermore, τ*_1_(*θ*) *is strictly decreasing and τ*_2_(*θ*)) *is strictly increasing for θ* ∈ (1*/*2, 1), *such that*

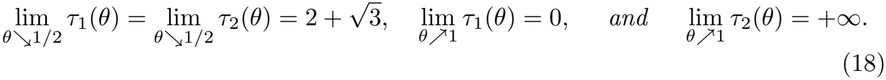

*Proof* First, we claim that max *f*_*ϵ*_(*x*) *>* 0 if and only if 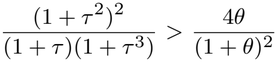 . To see this claim, notice that *f*_*ϵ*_(*x*) = *−*(1 + *τ* ^3^)*x*^3^ + (1 + *θ*)(1 + *τ* ^2^)*x*^2^ *−* (1 + *τ*)*θx* = *x*(*ax*^2^ + *bx* + *c*) where *a* = *−*(1 + *τ* ^3^) *<* 0, *b* = (1 + *θ*)(1 + *τ* ^2^) *>* 0 and *c* = *−*(1 + *τ*). Since 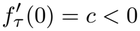 and, we deduce that 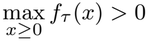 if and only if *f*_*ϵ*_has two positive roots, or equivalently, the quadratic equation *ax*^2^ + *bx* + *c* = 0 has two positive roots. Since *a <* 0 *< b*, this is equivalent to Δ = *b −* 4*ac >* 0, which is equivalent to 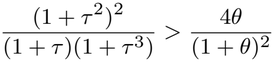 .

Next, we let

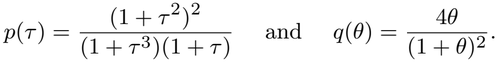

From calculus, it is easy to obtain

i. *p*(1) = 1, *p>*(+*∞*) = 1, *p*(*τ*) is strictly decreasing in 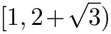 and strictly increasing in 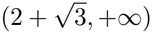 . Moreover, 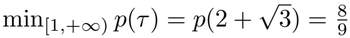 .
ii. *q* is strictly increasing in (0, 1), 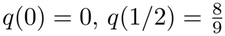 and *q*(1) = 1.

Therefore, for each *θ* ∈ (0, 1*/*2) and any *τ ≥* 1, we have 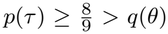 . Hence *p*(*τ*) *> q*(*θ*) and max_*ϵ*≥ 0_ *f*_*ϵ*_(*x*) *>* 0 by (i). Since 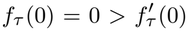, then *f*_*ϵ*_(*x*) has two additional positive roots *x*^*^(*τ*) *< x*^**^(*τ*). For the third order polynomial *f*_*ϵ*_to have three distinct roots, each of *x*^*^ and *x*^**^ must be non-degenerate and thus depend continuously on *τ* ∈ [1, +*∞*) (by the implicit function theorem). Finally, setting *τ* = 1, we have *f*_1_(*x*) = 2*x*(1 *− x*)(*x − θ*) and we see immediately that *x*^*^(1) = *θ* and *x*^**^(1) = 1. This proves part (a).

Next, we show part (b). For *θ* ∈ (1/2, 1), the fact that 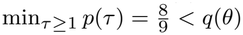, properties (iii) and (iv) above, and the Intermediate Value Theorem imply the existence of 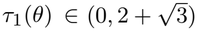 and 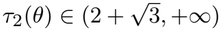 such that

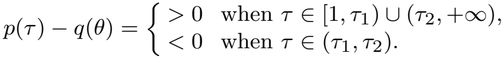

And differentiating *p*(*τ*_*ϵ*_(*θ*)) *− q*(*θ*) = 0 in *θ*, we obtain that 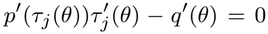 for *j* = 1, 2, so 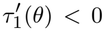 and 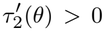 from properties (i) and (ii) above, and the limiting behaviors in (18) hold. Furthermore, for each *θ* ∈ (1*/*2, 1) and *τ* ∈ (*τ*_1_, *τ*_2_), *p*(*τ*) *− q*(*θ*) *<* 0 and hence *f*_*ϵ*_(*x*_*ϵ*_) *<* 0, which implies that *f*_*ϵ*_(*x*) *<* 0 for all *x >* 0. This proves part (b).

*Proof of Theorem 3*.*3* Fix *θ* ∈ (0, 1/2). If *d*_21_ = *d*_12_, then it is obvious that (*θ, θ*) and (1, 1) are two positive equilibria. Henceforth we may assume without loss of generality that *d*_12_ *> d*_21_ are given (otherwise relabel the patches). By Lemma 3.5(a), for every *τ >* 1, the function *f*_*ϵ*_has at least two roots 0 *< x*^*^(*τ*) *< x*^**^(*τ*) *<* 1. It remains to choose *τ* ∈ (1, *∞*) such that

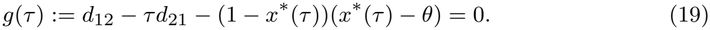

Now, *x*^*^ *→ θ* and *x*^**^ *→* 1 as *τ →* 1, so

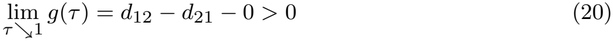

Also, *x*^*^, *x*^**^ remains bounded (within the interval [0, 1]) as *τ → ∞*, (in fact, both of them tend to zero) so

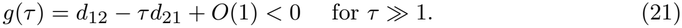

Hence, by the continuity of *g*(*τ*) and the Intermediate Value Theorem, there exists at least one *τ* ^*^ ∈ (1, ∞) so that *g*(*τ* ^*^) = 0, and we obtain one equilibrium (*x*^*^(*τ* ^*^), *τ* ^*^*x*^*^(*τ* ^*^)). Arguing similarly with *x*^**^(*τ*) in place of *x*^*^(*τ*), we obtain a second equilibrium. □

### 3.3 Nonexistence of positive equilibria

We first prove the following result on the bound of an equilibrium of (3).

#### Lemma 3.6.

*Let* (*x, y*) *be a nonnegative equilibrium of* (3), *and* (*x, y*) *≠* (0, 0). *If d*_12_ *≤ d*_21_ *and θ >* 0, *then* 0 *< x ≤ d*_21_*/d*_12_ *and* 0 *< y ≤* 1; *and similarly if d*_21_ *≤ d*_12_, *then* 0 *< x ≤* 1 *and* 0 *< y ≤ d*_12_*/d*_21_.

*Proof* If (*x, y*) is a nonnegative equilibrium of (3), then

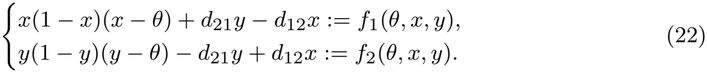

Without loss of generality, we only prove the case of *d*_12_ *≤ d*_21_. It is easy to see that *x* = 0 if and only if *y* = 0, since (*x, y*) *≠* (0, 0) then *x >* 0 and *y >* 0. Suppose that the result does not hold, then either *y >* 1 or *x > d*_21_*/d*_12_.

#### Case 1

*y >* 1. Then from the second equation of (22) we obtain

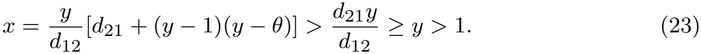

Then from the first equation of (22) and (23) we get

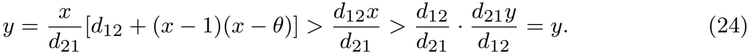

That is a contradiction. Hence *y ≤* 1.

Case 2: *x > d*_21_*/d*_12_. Then from the first equation of (22) we have

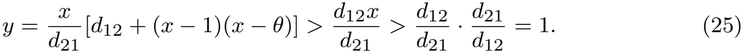

We can use the proof in Case 1 to get a contradiction again. Hence *x ≤ d*_21_*/d*_12_.

From Cases 1 and 2, we conclude that 0 *< x ≤ d*_21_*/d*_12_ and 0 *< y <* 1. □

Now we prove the following result about the nonexistence of positive equilibria of (3) when *θ* is greater than 1*/*2 and less than 1.

#### Theorem 3.7.

*Let θ* ∈ (1*/*2, 1) *and let* 1 *< τ*_1_(*θ*) *< τ*_2_(*θ*) *be given by Lemma 3.5. Then for each bounded interval K* ⊂ (1*/τ*_2_(*θ*), 1*/τ*_1_(*θ*)) ∪ (*τ*_1_(*θ*), *τ*_2_(*θ*)), *there exists E*_*K*_ *«* 1 *such that for all*

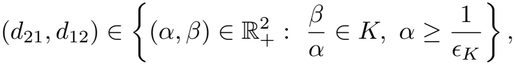

*the system* (3) *has only the trivial equilibrium* (0, 0). *Furthermore*, (0, 0) *is globally asymptotically stable*.

*Proof* Let *θ* ∈ (1/2, 1) and *K* be given as in the statement of the theorem. By Theorem 3.1 and Proposition 3.2, it remains to show the existence of some *E*_0_ *>* 0, such that for each

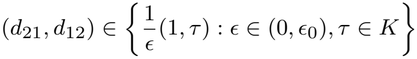

the system (12) has no positive equilibrium.

Suppose to the contrary that for a sequence, *E →* 0, the system (12) with 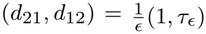 (where *τ*_*ϵ*_∈ *K*) has a positive equilibrium (*x*_*ϵ*_, *y*_*ϵ*_). Without loss of generality, we may assume that *τ*_*∞*_ := lim_*ϵ→*0_ *τ*_*ϵ*_*>* 1.

#### Claim 1.

*Either x*_*ϵ*_∈ (*θ*, 1) *or y*_*ϵ*_∈ (*θ*, 1).

Otherwise *x*_*ϵ*_(1 *− x*_*ϵ*_)(*x*_*ϵ*_*− θ*) + *y*_*ϵ*_(1 *− y*_*ϵ*_)(*y*_*ϵ*_*− θ*) *<* 0 which is a contradiction (for the sum of two equations of (12) should be zero).

#### Claim 2.

*There exists C ≥* 1 *independent of E such that* 0 *≤ x*_*ϵ*_*≤ C and* 0 *≤ y*_*ϵ*_*≤ C*.

By Claim 1, we may assume without loss that *x*_*ϵ*_∈ (*θ*, 1) for all *E «* 1, then (rewrite (12) using 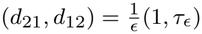 to get)

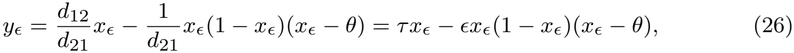

which is also bounded uniformly in *E «* 1.

#### Claim 3.

*Passing to a subsequence, there exists x*_*∞*_ *>* 0 *such that* (*x*_*ϵ*_, *y*_*ϵ*_) *→* (*x*_*∞*_, *τ*_*∞*_*x*_*∞*_).

By Claim 2, we may pass to a subsequence and assume that (*x*_*ϵ*_, *y*_*ϵ*_) *→* (*x*_*∞*_, *y*_*∞*_). By Claim 1, (*x*_∞_, *y*_∞_) ≠ (0, 0). To see Claim 3, let *E →* 0 in (26), and obtain

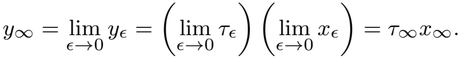

By Claim 1, at least one of *x*_*∞*_ or *y*_*∞*_ is positive and thus both are positive. Claim 3 follows.

#### Claim 4.

*Let x*_*∞*_ *>* 0 *be given in Claim 3, we must have* 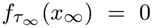 *and τ*_*∞*_ ∈ (*τ*_1_(*θ*), *τ*_2_(*θ*)).

By the first equation of (17), and Claim 3, we have

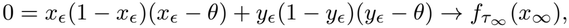

and *τ*_*∞*_ ∈ *K* ⊂ (1*/τ*_2_(*θ*), 1*/τ*_1_(*θ*)) ∪ (*τ*_1_(*θ*), *τ*_2_(*θ*)). Since *τ*_*∞*_ *>* 1, we have *τ*_*∞*_ ∈ (*τ*_1_(*θ*), *τ*_2_(*θ*)). Claim 4 is proved.

By Claim 4, 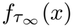 has at least one positive root. However, this is impossible in view of *τ*_*∞*_ ∈ (1*/τ*_2_(*θ*), 1*/τ*_1_(*θ*)) ∪ (*τ*_1_(*θ*), *τ*_2_(*θ*)) and the definition of *τ*_1_(*θ*), *τ*_2_(*θ*) in Lemma 3.5(b). This contradiction proves Theorem 3.7. □

Now we prove the following result about the nonexistence of positive equilibria of (3) when *θ* is sufficiently close to 1.

#### Theorem 3.8.

*For any d*_12_, *d*_21_ *>* 0 *and d*_12_ ≠ *d*_21_, *there exists* δ *>* 0 *such that when* 1 *−* δ *< θ <* 1, *the system* (3) *has only the trivial equilibrium* (0, 0) *which is globally asymptotically stable*.

*Proof* We first consider the system (3) in the case of *θ* = 1, that is:

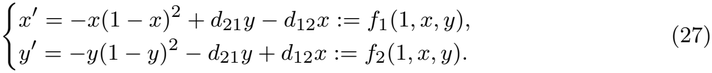

We claim that the nullclines *f*_1_(1, *x, y*) = 0 and *f*_2_(1, *x, y*) = 0 of (27) intersect only at (0, 0) in the closure of the first quadrant. Indeed, *f*_1_(1, *x, y*) = 0 and *f*_2_(1, *x, y*) = 0 can be rewritten as

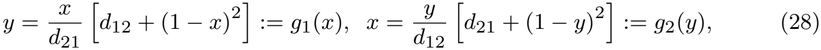

respectively. If (*x, y*) is an intersection point of the nullclines *f*_1_(1, *x, y*) = 0 and *f*_2_(1, *x, y*) = 0 such that *x ≥* 0 and *y ≥* 0, then from (28),

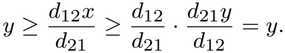

Hence we must have *d*_21_*y* = *d*_12_*x*, which implies *x*(1 *− x*)^2^ = 0 from *f*_1_(1, *x, y*) = 0, and *y*(1 *− y*)^2^ = 0 from *f*_2_(1, *x, y*) = 0. Therefore (*x, y*) = (0, 0), or (*x, y*) = (1, 1) and *d*_12_ = *d*_21_. Since *d*_12_ *≠ d*_21_, then (*x, y*) = (0, 0). This shows that (0, 0) is the only nonnegative equilibrium in 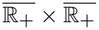 when *θ* = 1.

Now we consider the case that *θ* is near 1. Since *d*_12_ *≠ d*_21_, then without loss of generality, we assume that *d*_21_ *> d*_12_. From Lemma 3.6, a nonnegative equilibrium (*x, y*) satisfies 0 *≤ x ≤ d*_21_*/d*_12_ and 0 *≤ y ≤* 1. Let 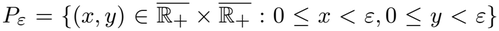 for some small *ε >* 0 . Then (0, 0) is an equilibrium in *P*_*ϵ*_for any *θ >* 0. But otherwise, if (*x, y*) ∈ *P*_*ϵ*_, (*x, y*) *≠* (0, 0) and *f*_1_(*θ, x, y*) = 0, then *y > d*_12_*x/d*_21_, while if (*x, y*) ∈ *P*_*ϵ*_, (*x, y*) *≠* (0, 0) and *f*_2_(*θ, x, y*) = 0, then *x > d*_21_*y/d*_12_. Hence (0, 0) is the only equilibrium in *P*_*ϵ*_. Therefore any other nonnegative equilibria of (27) for the case of *d*_21_ *> d*_12_ and *θ >* 0 is contained in the set

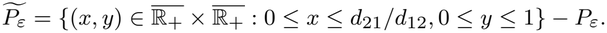

Now from the last paragraph, when *θ* = 1, there are no nonnegative equilibria of (3) in 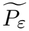 . That means, the curves 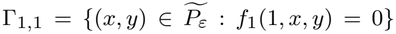 and 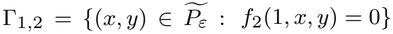 have no common points. Since 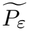 ;is a compact subset of R^2^, then the distance between Γ_1*ϵ*1_ and Γ_1*ϵ*2_ is positive, say *ξ >* 0. Since the functions *f*_1_(*θ, x, y*) and *f*_2_(*θ, x, y*) are continuous, then there exists δ *>* 0 such that the distance between Γ_*ϵ*1_ and Γ_*ϵ*2_ is greater than *ξ/*2 *>* 0, if 1 *−* δ *< θ <* 1 + δ, where 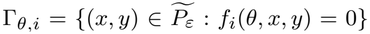 for *i* = 1, 2. This implies that there are no equilibria of (3) in 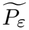 ;, and consequently, (0, 0) is the only nonnegative equilibrium of (3). □

The result in Theorem 3.8 has a clear biological meaning: if the dispersal rates *d*_12_ and *d*_21_ are unequal, then global extinction occurs if the Allee threshold is high enough. This can also be seen from the bifurcation diagram in Fig. 1: if (*d*_12_, *d*_21_) does not lie on the diagonal, then it will be in the global extinction region if the Allee threshold *θ* is close enough to 1. Indeed when *θ* is close to 1, the global extinction region almost covers the entire square {(*d*_12_, *d*_21_) : 0 *≤ d*_12_, *d*_21_ *≤* 1}. On the other hand, we note that the condition *d*_12_ *≠ d*_21_ is necessary for global extinction for any *θ* ∈ (0, 1), as when *d*_12_ = *d*_21_, there exist at least three nonnegative equilibria (0, 0), (*θ, θ*) and (1, 1).

**Fig 1.**
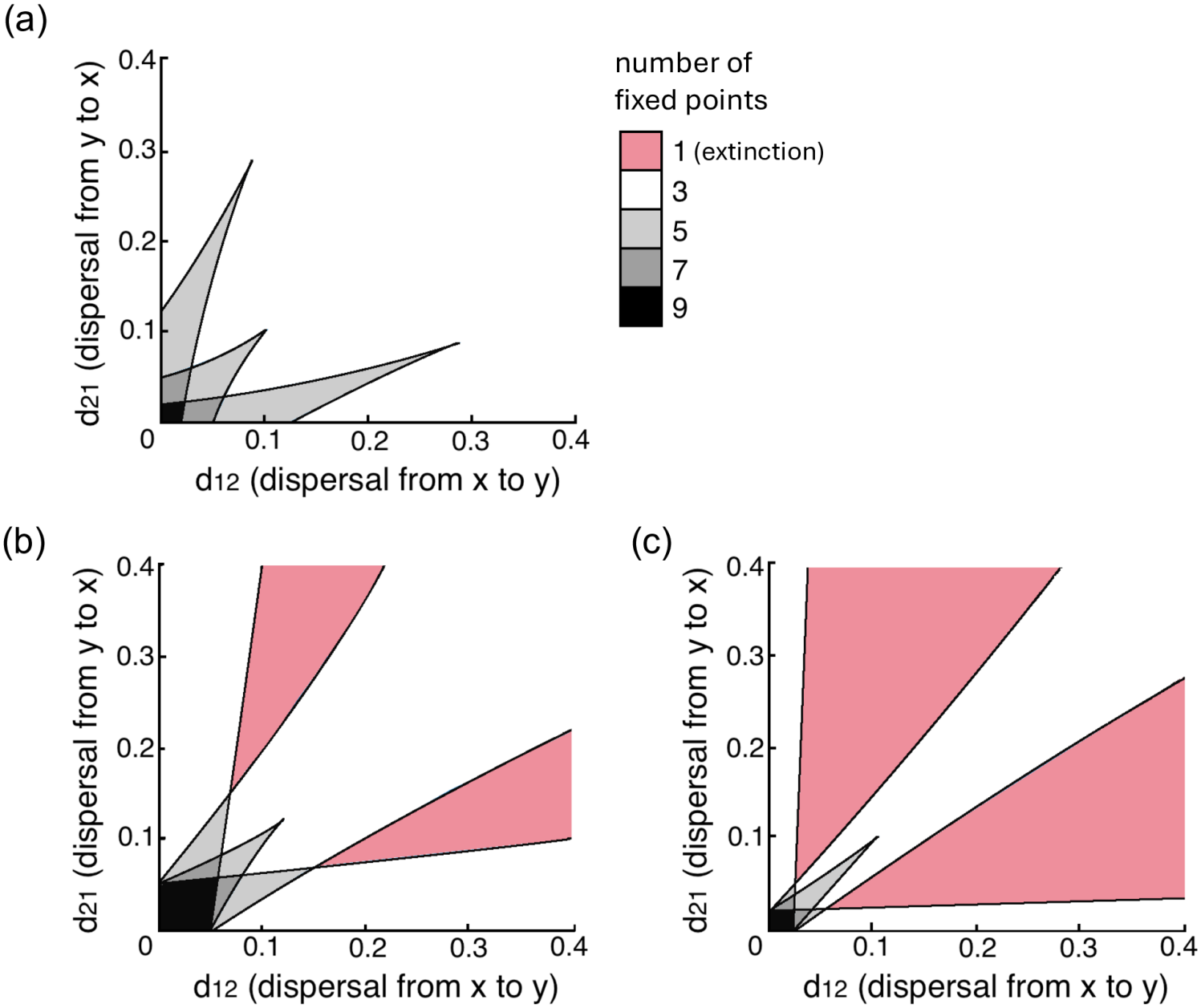
Two-dimensional bifurcation diagrams show the evolution of number of equilibria of system (3) when the Allee threshold *θ* increases past the critical separation value *θ*_*ϵ*_= 0.50 (shown in Theorem 3.3 and 3.7). At *θ* = 0.30, (a) is the typical appearance of the bifurcation diagram when *θ < θ*_*ϵ*_. Note the absence of any (pink) region where the extinction state (0, 0) is the sole equilibrium of the system. At *θ* = 0.55 and *θ* = 0.70, (b) and (c) show the encroachment of the extinction zone (in pink) as *θ* approaches 1.

## 4 Numerical Results

Numerical analysis of our two-patch asymmetric model (3) comes in three parts. We first create a two-dimensional bifurcation diagram as a general map of system behavior (Section 4.1). Second, we analyze one-dimensional bifurcations evident in the system, moving along our two-dimensional bifurcation diagram for three values of *θ* (Section 4.2). Last, our phase portraits, representative of the system’s behavior at noteworthy parameter values (Section 4.3).

The Matlab package `Matcont` (Dhooge et al. 2008) was instrumental in tracking solutions from noteworthy initial equilibrium points, using both *d*_12_ and *d*_21_ as bifurcation parameters for the two-dimensional diagram in Section 4.1, and *d*_21_ as the bifurcation parameter for Section 4.2 (keeping *d*_12_ fixed at 0.07). We then used the Matlab package `Pplane` (Arnold and Polking 1000) to create the phase portrait diagrams in Section 4.3.

### 4.1 Two-Dimensional Bifurcations

Theorems 3.3 and 3.7 reveal two distinct dynamical regimes for system (3), depending on whether the Allee threshold *θ* lies below or above the critical value *θ*_*s*_ = 0.50. For this reason, we open our numerical section by inspecting the bifurcation behavior of both dispersal parameters, at three separate values of the Allee threshold. Fig. 1 is a snapshot of number of equilibria of our system at these values of *θ*: Each shaded region of our diagram has a certain number of equilibria, and each boundary between regions is a bifurcation curve of equilibria where the number of equilibria change by two through a saddle-node or pitchfork bifurcation.

When *θ < θ*_*s*_ = 0.50, as in Fig. 1(a), there is always at least one nontrivial, stable equilibrium for (3). In context of our motivation, this means that for Allee thresholds below 0.5, there is always a set of populations for *x* and *y* that is is sustained above extinction. When the values of *d*_12_ and *d*_21_ are very small (*<* 0.01), there are 9 equilibria (as was seen by Kang and Lanchier (2011) for symmetric small dispersal). As we increase *d*_12_ and *d*_21_ along the diagonal line *d*_12_ = *d*_21_, two pairs of fixed points simultaneously collide in saddle-node bifurcations, leaving us with only five fixed points. This happens once more—in Fig. 1(a), at approximately (0.1, 0.1)—with the system maintaining three equilibria for the remainder of the dispersal parameter values. Note how the system’s behavior at or around the line *d*_12_ = *d*_21_ does not qualitatively change even when *θ > θ*_*c*_ in our successive diagrams. This is why, in previous work with symmetrical systems, our novel extinction state was not evident (Kang and Lanchier 2011).

When *θ* passes *θ*_*s*_ = 0.50, as in Fig. 1(b) and (c), areas containing only one the trivial equilibrium (0, 0) appear. These extinction zones steadily expand with larger values of *θ*. In these ranges of (*d*_12_, *d*_21_) parameter values, populations in both patches are depleted and cannot recover, no matter the initial population. Recall that our model (3) has been scaled to have a population limit of 1 for each patch; as the Allee threshold approaches 1 it becomes more and more difficult for the populations *x* and *y* to sustain either themselves or each other.These extinction regions of the dispersal rates are concerning for biological conservation efforts; fortunately, although our model can predict these effects, it is unrealistic to expect that a population would have an Allee threshold as high as *θ*_*s*_ = 0.5.

### 4.2 One-Dimensional Bifurcations

One-dimensional bifurcation diagrams with only one dispersal rate now allow us to focus on qualitative changes within a small section of our larger two-dimensional diagram. Of especial interest are those red regions from Fig. 1(b) and (c) which promote inevitable extinction: How do they appear, and what does the system look like at that instant?

For context, imagine fixing a vertical line at *d*_12_ = 0.07 on any of the three diagrams in Fig. 1. We created the bifurcation diagrams in Fig. 2 by moving a point upwards along this line from *d*_21_ = 0 to *d*_21_ = 0.25 for three different values of *θ*. The appearance and disappearance of positive equilibria due to parameter variation is indicated in Fig. 1 as the curves between differently shaded sections, and in Fig. 2 as a coming-together or birth of two equilibria branches (colored curves) in a saddle-node or pitch-fork bifurcation. This connection between the two diagrams is useful to remain oriented within the framework of Section 4.1.

**Fig 2.**
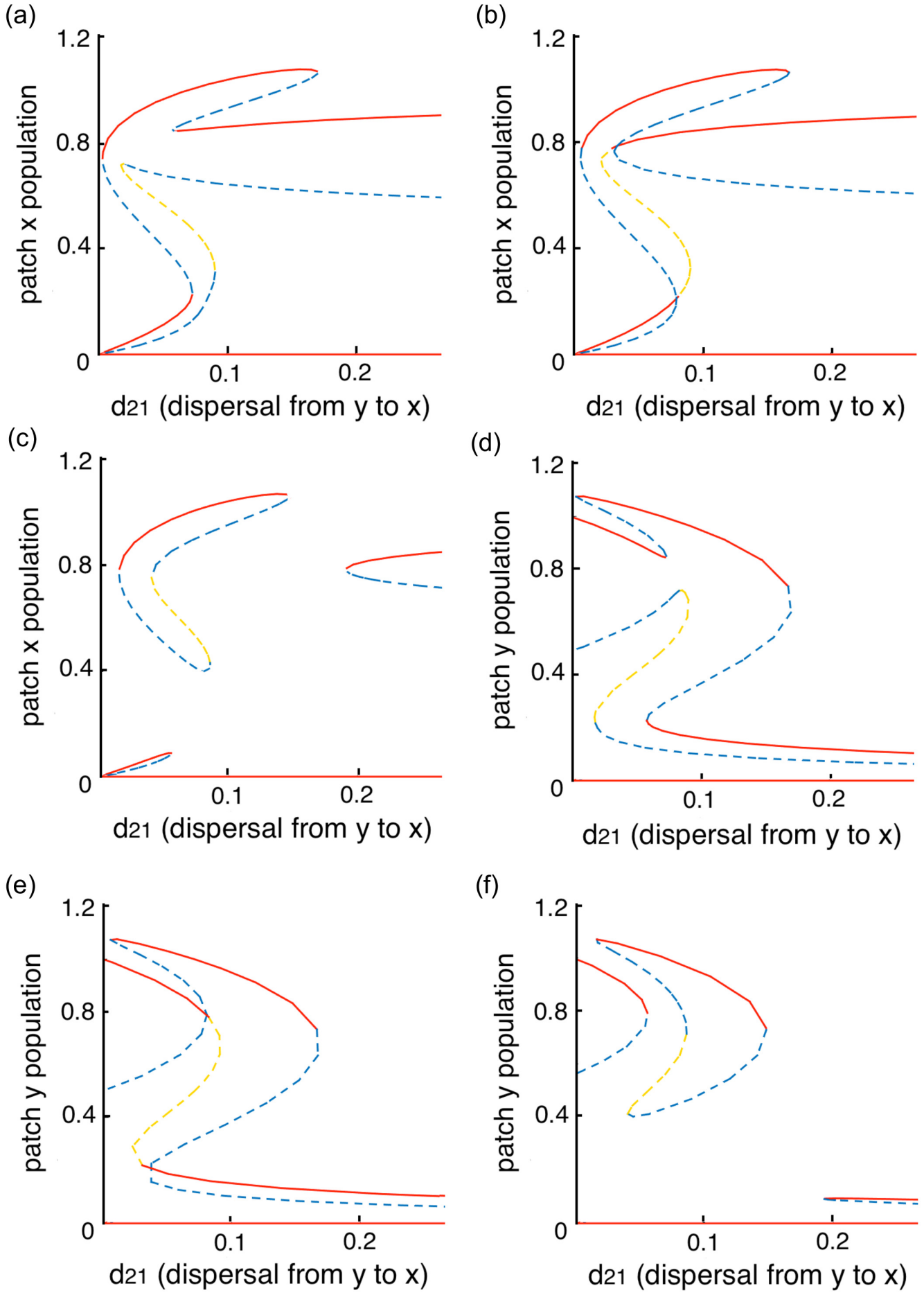
A one-dimensional bifurcation diagram with bifurcating parameter *d*_21_, fixed value *d*_12_ = 0.07, and varying values of *θ*. The left-hand column represents the population of patch *x*; the right column, patch *y*. Red curves indicate trajectories of stable equilibria; blue, saddle points; and yellow, the source. For (a) *θ* = 0.49 *< θ*_*ϵ*_; (b) *θ* = *θ*_*ϵ*_= 0.50; and (c) *θ* = 0.56 *> θ*_*ϵ*_. The critical value *θ*_*ϵ*_= 0.50 is the imperfect bifurcation point at which an isola separates.

Fig. 2 shows the progression of our system from a *θ* value below the separation point (*θ < θ*_*s*_), at the transition (*θ* = *θ*_*s*_ = 0.50), and afterwards (*θ > θ*_*s*_), for both the *x* (Fig. 2, left column) and *y* (Fig. 2, right column) patches. Each curve is a branch of positive equilibria, from birth to death. We chose a small value for *d*_12_ = 0.07 in order to show a variety of structures; low dispersal values showcase the most diverse number of states for system (3).

When *θ* = 0.50 (Fig. 2 (b,e)), the set of positive equilibria is a connected set in (*d*_21_, *x, y*) space, and there are four saddle-node bifurcation points (two forward and two backward) and two pitchfork (one forward and one backward) bifurcation points. With *d*_21_ increasing from zero, three forward bifurcations occurs and two of them generate a new stable positive equilibrium, then three backward bifurcation brings the number of equilibria back to 3 (which is the number for sufficiently small *d*_21_).

For *θ* = 0.49 (a small perturbation from *θ* = 0.50), the bifurcation diagram changes dramatically from the one for *θ* = 0.50 (Fig. 2 (a,d)). The set of positive equilibria is no longer a connected set in (*d*_21_, *x, y*) space, and instead it breaks up into two connected components. Now there are six saddle-node bifurcation points (three forward and three backward). The breakup of a one-dimensional pitchfork bifurcation into two separate branches is known as an imperfect pitchfork bifurcation for a two-dimensional bifurcation problem (Liu et al. 2007) (see also Ruan and Xiao (2023); Shi (1999)).

One distinct characteristic of our system (3) is the appearance of isolas in the one-dimensional bifurcation diagram. An isola represents a connected component of the equilibrium set that is separate from the main connected component, forming a mushroom-like shape (Gray and Scott 1985). The bifurcation diagram for *θ* = 0.56 (Fig. 2(c, f)) indicates the existence of an isola in system (3). The creation of the isola shows the behavior of the imperfect pitchfork bifurcation for *θ > θ*_*s*_ = 0.50 (the critical value for parameter *θ*). Similar to the case *θ* = 0.49 before the critical value, there are six saddle-node bifurcation points (three forward and three backward), but they appear in an order of FFBBBF, while they are in an order of FFFBBB when *θ < θ*_*s*_. In the gap between the final backward and forward saddle-node bifurcation points, there is only one equilibrium, the trivial one (0, 0), which corresponds to the extinction region in the two-dimensional diagram Fig. 1(b,c). We remark that if *θ* takes a larger value or *d*_12_ is larger, there could be two extinction gaps for *d*_21_ bifurcation diagram in which the six saddle-node bifurcations occur in an order of BFFBBF. A segment BF in the sequence of saddle-node bifurcations is the signature of an extinction parameter region.

When there are two extinction gaps in *d*_21_ bifurcation diagram (for large *θ*), the set of positive equilibria consists of three connected components. The three persistence parameter regions correspond to the cases when the ratio of the dispersal rates *d*_12_*/d*_21_ is either sufficiently large, sufficiently close to 1, or sufficiently small. The isola is the case of *d*_12_*/d*_21_ is sufficiently close to 1. That means, populations in two patches can achieve coexistence when they take the same dispersal rate (strategy), but if one population (say, the one in patch 2) either increases or decreases the dispersal rate continuously, then a global extinction will occur in either scenario. The only safe strategy for population in patch 2 not going extinct will be staying within the parameter interval of that isola.

### 4.3 Phase Portraits and Basins of Attraction

We now move from bifurcation analysis towards the use of phase portraits, to focus on the individual steady states of our *x* and *y* patches for different parameter values. To place Fig. 3 in context, refer to Fig. 1(b). Again imagine a vertical line at *d*_12_ = 0.04 and follow the transition of the system through the different states as *d*_21_ increases.

**Fig 3.**
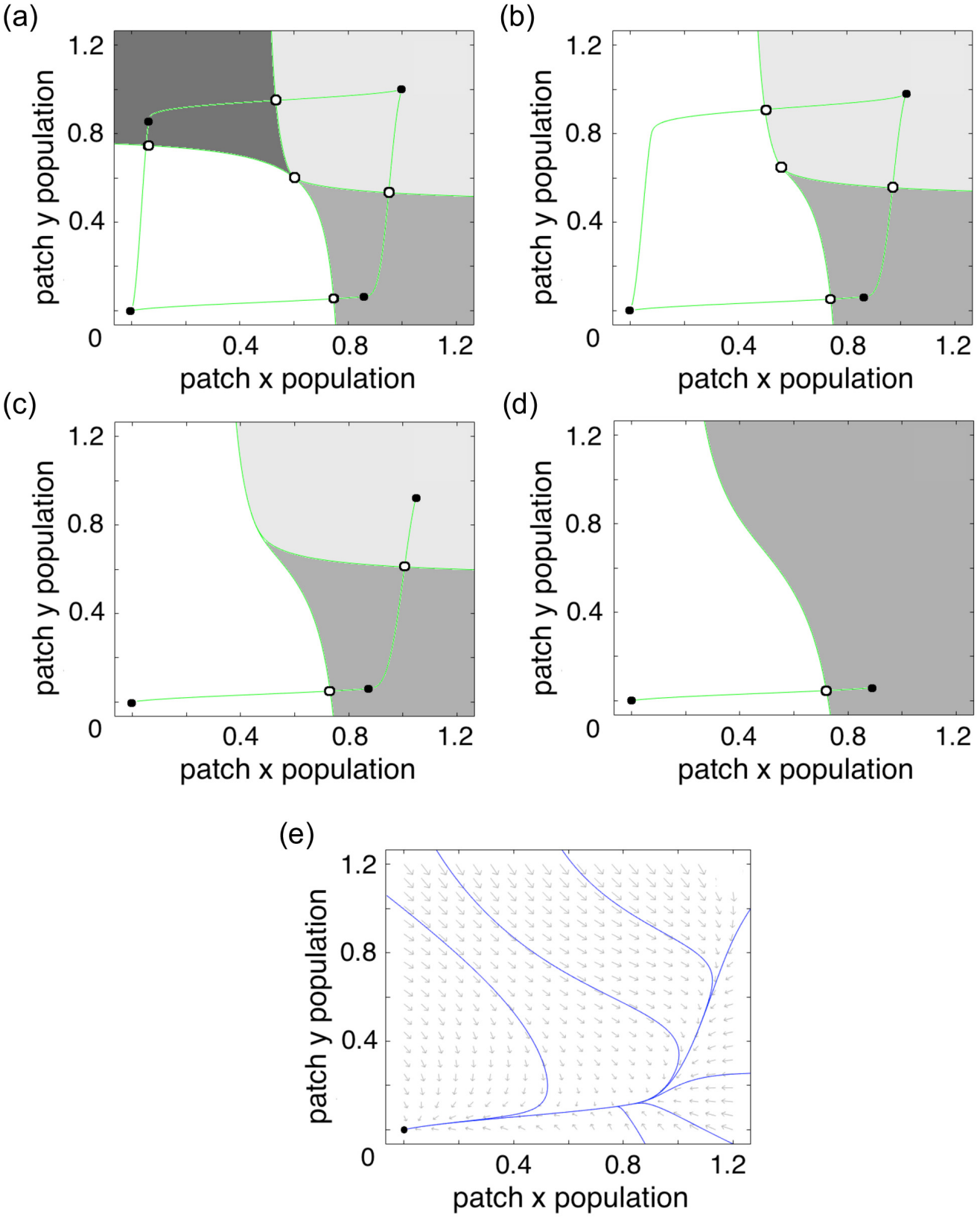
Basins of attraction of stable equilibria for varying values of *d*_21_. Here the Allee threshold (*θ* = 0.55) and dispersal from patch *x* to *y* (*d*_12_ = 0.04) are constant throughout each panel, whereas *d*_21_ increases for each (0.04, 0.06, 0.07, and 0.10 for (a-d)). Each solid point is a stable equilibrium, while hollow points are unstable equilibria. Basins of attraction are distinguished by varying shades of gray. The green curves connecting equilibria are the stable and unstable manifold for each saddle point, connected to the unstable source in the center. Each successive phase portrait is created by a saddle-node collision of two equilibria points in the previous diagram. Panel (e) shows the phase portrait for *d*_21_ = 0.11: there is only the trivial equilibrium and it is globally asymptotically stable.

In Fig. 3(a), the symmetric case, there are 4 stable equilibria (9 total); this case is well-known and covered in detail by Kang and Lanchier (2011). Moving from Fig. 3(a) to (b), we are presented with the first of two saddle-node bifurcations. The equilibria with high *y* and low *x* has collided with its neighboring saddle point, and has disappeared in a saddle-node bifurcation. Fig. 3(c) and (d) are created in the same way, until we are left with only one nontrivial sink. This is in keeping with our initial observation, as now the only stable equilibrium occurs at high values of the *x* patch’s population. In a real-world context, this makes perfect sense: if the dispersal from patch *x* is kept at a very low value, and the dispersal from patch *y* increases, eventually patch *y* will be depleted and patch *x* will have a much better chance at survival. When *d*_21_ is increased to 0.11 in Fig. 3(e), the final nontrivial equilibrium also vanishes, population in patch *y* drops first, and population in patch *x* decreases to zero as well.

Fig. 4 shows the progression of nullclines as these bifurcations occur. We chose slightly different values of our dispersal parameters in order to highlight the appearance and disappearance of the extinction region mentioned in Section 4.2. As of now, we have not discovered a sensible biological reasoning behind the disappearance and reappearance of the second and third equilibria from Fig. 4(b-d), but we can make some observations. Firstly, that in this state, the trivial equilibrium has a larger basin of attraction, and the equilibria are more clustered, than in Fig. 3(b). Secondly, the two equilibria that reappear after the extinction zone are not the same two that disappeared in the saddle-node bifurcation from Fig. 4(b-c); these two are at a comparable value of *x*, but a much lower value of *y*; this suggests that, while the population of *y* can be recovered with sufficiently high dispersal from *y* to *x* (again, biological reasoning for this phenomenon is lacking), its stable population is very much smaller than before.

**Fig 4.**
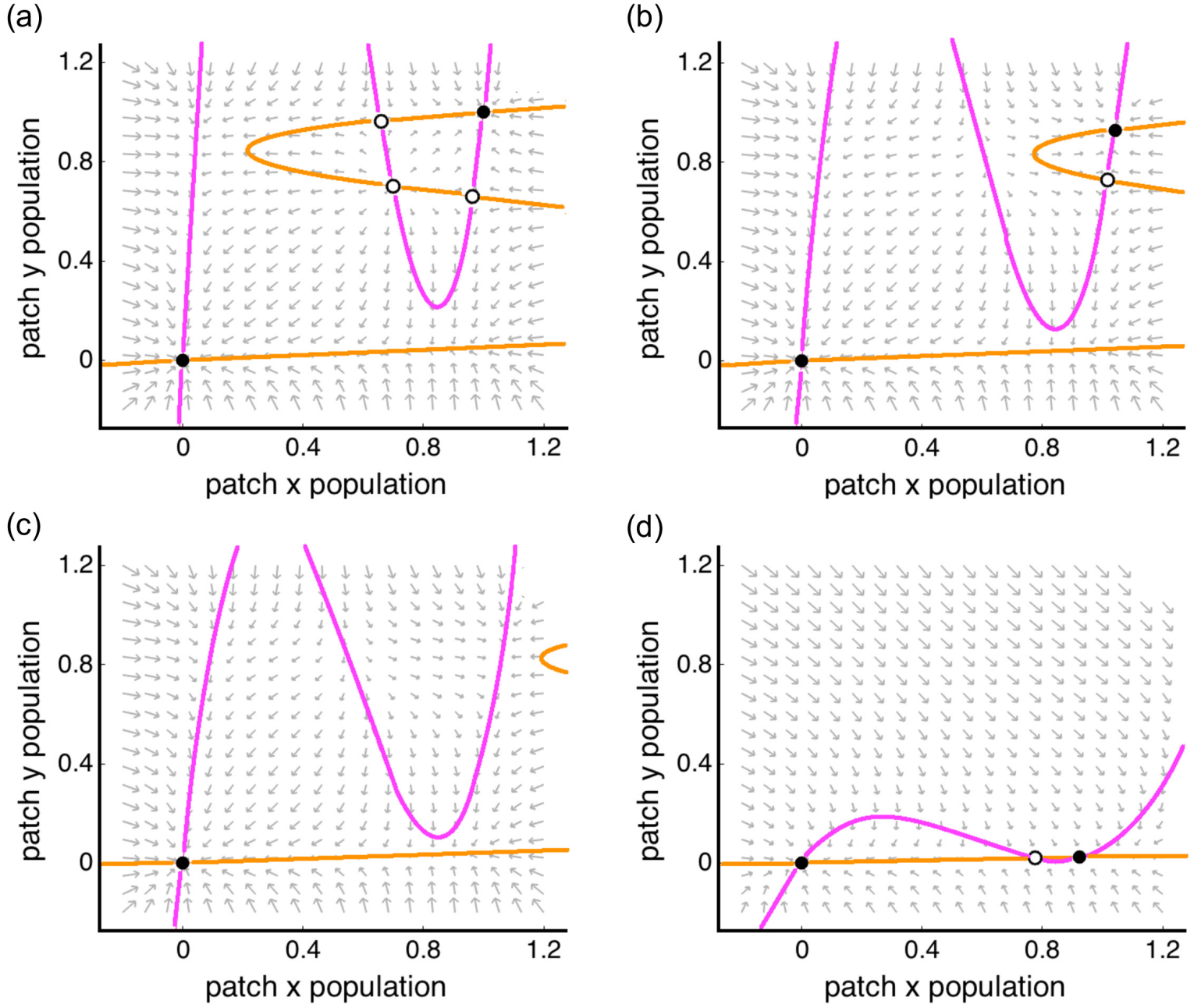
A phase portrait showing the system’s nullclines. *θ* = 0.70 and *d*_12_ = 0.03 are held constant, whereas *d*_21_ increases for each panel: 0.03, 0.05, 0.065, and 0.50 for (a-d). Solid points are stable equilibria and hollow points are unstable ones. The *x* nullclines are represented by the color pink, and *y* with orange. From (a) (symmetric) to (b), the two leftmost unstable equilibria collide; from (b) to (c) the two in the northeast corner. From (c) to (d), however, there is a creation of two more fixed points, a seeming reversal of the previous bifurcation, but at a much lower population of *y*.

## 5 Discussion

Here, we investigated the rich dynamics of a two-patch metapopulation model with asymmetric dispersal and an Allee effect. To represent the Allee-effect growth dynamics, we employed the simplest cubic growth function as a prototype for a broader class of nonlinear growth functions. The corresponding single-patch model exhibited a well-known bistable structure, with extinction and persistence as the two stable states and the Allee threshold separating their basins of attraction. For the coupled two-patch system, we showed that every solution converges to an equilibrium in long time, ruling out sustained oscillatory behavior. The system can possess between one and nine non-negative equilibria, with their number and configuration depending sensitively on the dispersal parameters. Finally, we characterized how asymmetric dispersal in this system can enable a “rescue effect” in some parameter ranges, where populations below an Allee threshold may persist due to disperal from a high-population coupled patch of the same species.

In our two-patch model with an Allee effect, patch rescue occurs when the ratio of the two dispersal rates is sufficiently large, regardless of the Allee threshold, or when the Allee threshold is less than half of the carrying capacity. In the former case, strong dispersal asymmetry allows the patch with a high emigration rate, which would otherwise decline toward extinction because its initial population is below the Allee threshold and individuals are lost through outward dispersal, to be continuously replenished by immigration from the other, more robust patch with a larger initial population. This rescue mechanism has been investigated in previous studies, including two-patch models with mortality during dispersal (Amarasekare 1998), and is ecologically important because immigration can sustain populations that would otherwise become depleted, although their equilibrium population size may remain below the carrying capacity that could be achieved in the absence of dispersal. A similar rescue effect occurs when the Allee threshold is sufficiently low relative to the carrying capacity, in which case populations in both patches can be maintained above the extinction threshold through dispersal. This is particularly important in systems where patch persistence is critical, rather than total population size, as patch rescuing can lead to lower overall population (Grumbach and Hilker 2025).

The Allee threshold *θ* plays an important role in determining the number of equilibria of the system. When the Allee threshold *θ* is below half of the carrying capacity, *i*.*e*., 0 *< θ <* 1*/*2, system (3) always possesses at least two positive equilibria. Consequently, the population persists for sufficiently large initial populations, and the rescue effect occurs regardless of the dispersal rates *d*_12_ and *d*_21_. On the other hand, when the Allee threshold *θ* exceeds half of the carrying capacity, *i*.*e*., 1*/*2 *< θ <* 1, positive equilibria of (3) exist only when the ratio of the dispersal rates *d*_12_*/d*_21_ is either sufficiently large, sufficiently small, or sufficiently close to 1. In contrast, when this ratio lies in an intermediate range, system (3) admits no positive equilibria. In this regime, the population undergoes global extinction, and the rescue effect completely fails.

That is, despite full population conservation during dispersal, global extinction nevertheless occurs for certain combinations of asymmetric dispersal rates and a relatively high Allee threshold. These extinction regions arise from an interplay between a strong Allee effect and intermediate levels of dispersal asymmetry: the dispersal rates are neither sufficiently small to permit the coexistence of multiple positive equilibria nor sufficiently symmetric to maintain a balanced distribution of individuals between the patches, and they are also not sufficiently asymmetric to generate a rescue effect. Interestingly, when the two patches are completely disconnected, each population persists provided its initial size is sufficiently large. However, when the ratio of the dispersal rates falls within certain intermediate ranges, the two populations can sequentially decline below the Allee threshold and ultimately collapse, even when both patches initially contain populations well above the threshold. Thus, while dispersal and connectivity are generally recognized as important mechanisms for promoting the persistence of fragmented populations through demographic rescue and recolonization, our results demonstrate that, in the presence of a strong Allee effect, population-conserving dispersal can instead induce global extinction.

Future work could extend this two-patch model to systems with multiple patches or interacting species. As demonstrated in this paper, coupling two populations subject to the Allee effect already leads to substantially richer dynamics than those of an isolated patch. It is therefore natural to investigate how these dynamics evolve in three- or *n*-patch systems with asymmetric, population-conserving dispersal. For sufficiently small dispersal rates, an *n*-patch system may admit as many as 3^*n*^ non-negative equilibria, reflecting the multiple persistence and extinction states associated with the Allee effect. However, it remains an open question whether every solution of the *n*-patch system necessarily converges to an equilibrium, as is known for *n*-patch logistic systems, or whether genuinely oscillatory dynamics, such as periodic orbits, can emerge when three or more patches are coupled. Results in this paper are based on the assumption that local dynamics in the two patches are identical. Models incorporating stochastic simulations such as in (Vortkamp et al. 2020) or introducing multiple species as in (Sen and Sinha 2022) could reveal how demographic or environmental fluctuations influence the likelihood of extinction near bifurcation boundaries and could also characterize the trajectories of extinction.

## 6 Conclusion

Directional dispersal, whether through wind, water, or other ecological gradients, can fundamentally help or hinder population persistence in systems subject to the Allee effect. In the conservation and restoration of some endangered species, understanding how directed, asymmetric dispersal interacts with density-dependent processes can be crucial for the sustainable establishment of new population patches, the maintenance of existing patches, and the health of the species as a whole. By providing analytical and numerical analyses of critical parameter regions for global extinction and patch rescue, this work adds key mathematical insights into dispersal-mediated Allee effects in metapopulation systems.

## Declarations

### Data availability

No data were used for the research described in the article.

## Acknowledgements

M. Swift would like to thank the NSF-EXTREEMS QED Research Experience for Undergraduates program and the Department of Mathematics at William & Mary for their support for the work underlying this paper. Sarah Day at W&M deserves special thanks for encouraging M. Swift to pursue this REU, which started her research career.

